# Multiplex PCR reveals population structure in an inbred communal bird

**DOI:** 10.1101/2024.01.29.577869

**Authors:** Sarah Babaei, Leanne A. Grieves, Ben Evans, James S. Quinn

## Abstract

We sampled communally breeding pūkeko (*Porphyrio melanotus melanotus*, family Rallidae) populations on the North (Tāwharanui Park) and South (Otokia Reserve) Islands of New Zealand that differ in climate and ecology. North Island populations have year-round territories, philopatry, and form kin groups, resulting in inbreeding. South Island populations have seasonal territories, high dispersal rates, and form non-kin groups, leading to outbreeding. Given behavioural evidence of inbreeding we predicted that the North Island population would exhibit lower heterozygosity and higher inbreeding coefficients than the South Island population. We hypothesized that the South Island population originated via a range expansion from the north and predicted that South Island birds would have lower allelic diversity due to founder effects. To test these predictions, we developed microsatellite primers, optimized multiplex PCRs, and genotyped breeding groups from the North and South Island. Breeding groups from North Island were genetically differentiated, whereas population structure was not detected in the South Island birds. North Island birds had higher inbreeding coefficients and levels of within group kinship, but not allelic diversity, compared to South Island birds. Our results are thus inconclusive about whether the South Island population originated via a range expansion from the north. This pilot study validated microsatellite markers and PCR methods and is the first genetic analysis of population structure and relatedness within communal breeding pūkeko groups. These genetic tools will be used for larger-scale studies to help resolve the origins of the South Island population and provide further insights into the effects of ecology and behaviour on inbreeding, reproductive success, and population demographics in this species. Pūkeko may provide an excellent model for experimental analyses of inbreeding effects on wild avian populations without the attendant concerns that come with small, endangered populations. This work may thus inform conservation efforts, including translocations of endangered species.

## Introduction

Most endangered species have small and declining populations (Wilcove et al. 1992), which may lead to inbreeding. In some cases, this results in inbreeding depression and may contribute to the extinction vortex (Blomqvist et al. 2010), which is when the reduction in population size increases inbreeding depression, resulting in further population decreases and continued inbreeding (Brook et al. 2002). Ideally, endangered species are not subject to research efforts that negatively affect the study subjects. Instead, species in large and growing populations that naturally engage in inbreeding can be used to investigate the effects of inbreeding and inbreeding depression on population size and persistence (Kardos et al. 2016), which then may inform endangered species conservation efforts.

Inbreeding decreases heterozygosity and may lead to inbreeding depression (Keller and Waller 2002). Inbreeding depression is thought to stem from two sources. The first is that fitness declines when deleterious recessive alleles are more frequently homozygous because of inbreeding (Keller and Waller 2002). The second is that fitness may decline when advantageous heterozygous genotypes are less prevalent.

Species may avoid inbreeding through natal dispersal, sometimes by only one sex. This reduces risks of inbreeding if breeding occurs after dispersal (Pusey and Wolf 1996). Nonetheless, inbreeding is still observed in healthy wild populations despite potential negative effects (Keller and Arcese 1998; Langen et al. 2011). One potential benefit of inbreeding is that it may facilitate the purging of deleterious recessive alleles that would otherwise be less susceptible to removal in an outbred population because they would often be heterozygous with a wild type allele (Hedrick 1994; Keller and Waller 2002). However, selection against deleterious recessive alleles also decreases population size, which itself is a factor that can increase inbreeding (Langen et al. 2011).

Interestingly, a recent study found that animals rarely avoid mating with kin and identified publication bias favoring papers showing inbreeding avoidance (de Boer et al. 2021). Under conditions in which inbreeding is associated with purging of deleterious recessive genes, there may be indirect fitness benefits from inbreeding because of increased kin-shared alleles (Langen et al. 2011). More generally, the effects of inbreeding may vary over time, and among species and ecological contexts, and why, when, and with whom inbreeding occurs is not always apparent (Keller and Waller 2002). Thus, it is important to study inbreeding and its consequences in a diversity of organisms and contexts. This is especially important if studies of non-endangered species have the potential to inform our understanding of inbreeding depression in small populations of endangered species.

Pūkeko *(Porphyrio melanotus melanotus*, family Ralliade) are communally breeding birds found in eastern Australia and New Zealand. In New Zealand, pūkeko exhibit cooperative, polygynandrous breeding and joint laying, where members of a breeding group share a communal nest that multiple females lay eggs into, and breeding individuals mate with multiple breeding partners (Craig 1980; Jamieson 1997). We studied pūkeko from New Zealand’s North Island (Tāwharanui Regional Park) and South Island (Otokia Wildlife Reserve; Fig 1). The North Island population experiences a milder climate compared to the South Island population, allowing them to defend year-round territories (Jamieson 1997). North Island birds have low natal dispersal (Jamieson 1997), likely due to habitat saturation that limits dispersal options. Stable groups include a few breeding males, one to three breeding females, and non-breeding helpers (Jamieson 1997). Observational data (collected at Shakespear Park in the North Island; Fig 1) indicate that a combination of year-round territoriality and philopatry has led to inbreeding, often between first-degree relatives (Craig and Jamieson 1988), and high reproductive skew in North Island populations (Jamieson 1997). By contrast, birds in the South Island population experience a harsher climate and changes in food availability in the winter, forcing them to abandon territories annually (Jamieson 1997). During the non-breeding season, South Island birds join large foraging flocks and re-establish breeding groups in new configurations of group members each breeding season, without helpers, resulting in non-kin breeding groups that lack inbreeding (Jamieson 1997). South Island group sizes are smaller with one to three males, one or two females, and no helpers (Jamieson 1997).

**Fig 1.**
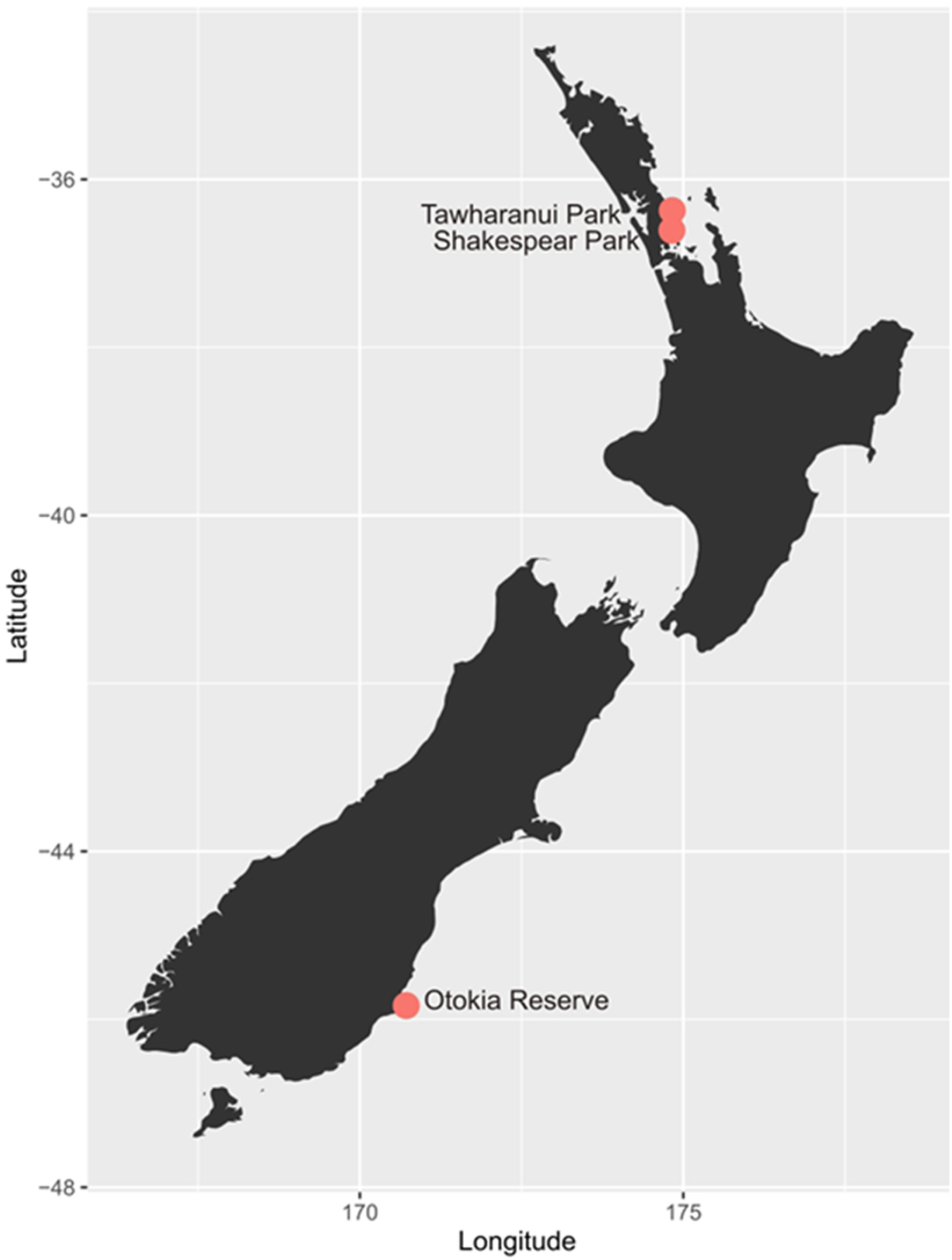
Map of New Zealand showing locations of North Island populations (Tāwharanui and Shakespear) and a South Island population (Otokia). Figure generated using *ggplot2* and *mapdata* R packages (Wickham 2016; Becker and Wilks 2018).

Overall, a high degree of kinship and inbreeding has been documented by demographic and behavioural observations in North Island populations (Craig and Jamieson 1988). The fitness consequences of this, however, are still unknown and there are limited genetic studies of the species. It is important to understand the degree to which inbreeding in the North Island populations has reduced within group heterozygosity and to what extent this causes inbreeding depression. This can be achieved by comparing North Island and South Island populations.

Pūkeko are listed as ‘Not Threatened’ by the International Union for Conservation of Nature (New Zealand Department of Conservation), making them tractable to study. The contrast between the North Island population, with nonbreeding helpers on year-round territories, and the South Island population, with outbred breeding groups on seasonal breeding territories without helpers, makes pūkeko a compelling model for exploring effects of inbreeding and kinship on inclusive fitness and population stability. To that end, we developed molecular tools for pūkeko. Our primary goal was to develop microsatellite markers and optimize multiplex PCR protocols that will facilitate large scale population genetic analyses in this study system. Additionally, because of the polygynandrous mating system and the likelihood that some matings involve non-kin, we anticipate future research opportunities to monitor the fate of inbred and outbred eggs in the same joint nest, enabling us to measure the effects of inbreeding on growth, survival, and fitness.

Previous genetic analyses of pūkeko used resource intensive techniques (Southern blotting) that were of lower genetic resolution and were difficult to assess statistically (Jamieson et al. 1994; Lambert et al. 1994). Here we describe the development of 18 polymorphic loci and a pilot study in which we compared inbreeding coefficients between the North Island and South Island populations, testing for genetic differentiation among breeding groups within populations. Based on prior evidence of inbreeding in a North Island population at Shakespear Regional Park located about 27 km south of Tāwharanui Regional Park (Craig and Jamieson 1988; Fig 1) and seasonal mixing of breeding groups in the South Island population (Jamieson 1997), we predicted higher inbreeding coefficients and greater within breeding group kinship in North Island (Tāwharanui) compared to South Island (Otokia). We predicted significant genetic structure based on breeding group differentiation within the North Island, but not South Island, based on a lack of local migration or membership exchange among breeding groups in the north. Finally, we also hypothesized that pūkeko from the North Island population historically expanded their range southward, leading us to predict lower allelic diversity in the South Island than the North Island, due to a founder effect.

## Material and Methods

### Sample Collection

This project used blood samples collected as part of long-term research on pūkekos. We selected four breeding groups from the North Island population sampled in 2010 by Dr. Cody Dey and J.S.Q. and two breeding groups from the South Island sampled in 1991 and 1992 by I. Jamieson (Jamieson et al.1994). For this pilot study we opportunistically selected larger breeding groups for which we had higher quality DNA. Individuals and breeding groups selected for each population are provided in S1 Table.

### Microsatellite Identification

Using one third of a lane of an Illumina 2500 machine, we collected short read data (150 base pair (bp) paired-end sequences) from three female pūkeko with barcoded DNA (one each from Tāwharanui park [North Island], Shakespear park [North Island], and Otokia [South Island]), prepared by The Centre for Applied Genomics. We used PEAR (Zhang et al. 2014) to merge paired end sequences, and MISA (Thiel et al. 2003; Beier et al. 2017) plus a Perl script to identify tetrameric microsatellites with at least 40 bp flanking sequence on each side of the repeat for primers. These data have been deposited in the NCBI Sequence Read Archive (BioProject doi: pending).

### Primer Design, Optimization and Genotyping

Primers were designed in Primer3 using tetramers with at least eight repeats (Koressaar and Remm 2007; Untergasser et al. 2012; Koressaar et al. 2018) based on the Illumina paired-end sequences described above. For multiplex PCR, which involves amplifying multiple loci in the same reaction, and subsequent genotyping, each forward primer was labeled at the 5’ end with fluorescent labels (HEX and 6FAM, Integrated DNA Technologies; NED, Applied Biosystems). PCR was performed in 25 µL reaction volumes containing nuclease-free water, 1X PCR reaction buffer, 2 mM MgCl_2_, 0.2 mM dNTPs (Invitrogen by ThermoFisher Scientific), 0.2 µM of each forward and reverse primer pair, 0.5 U Taq polymerase (ThermoFisher Scientific), and 1 µL template DNA (extracted from pūkeko blood samples using a standard salt extraction procedure; Miller et al. 1988). We used a thermocycling program of 3 min initial denaturation at 95°C, followed by 30 cycles of 30 sec denaturation at 95°C, 30 sec annealing at (50-60°C), 45 sec extensions at 72°C, and a 10 min final extension at 72°C. All reactions were performed using an MJ Research PTC-200 Thermocycler. Where possible, individual primer pairs were multiplexed in groups of three.

After PCR, we mixed 5 µL of PCR product with 1 µL of 6X loading dye (ThermoFisher Scientific) and ran samples on 2% agarose gels stained with RedSafe (FroggaBio) at 95V for 30 min and visualized under UV light (Bio-Rad ChemiDoc Imaging System, Image Lab) to confirm amplification at the expected allele size. PCR products were then loaded into 96-well plates, cleaned using an in-house ethanol precipitation PCR cleanup protocol (S2 File), and sent for genotyping on a 48-capillary ABI 3730 at the Trent University Wildlife Forensic DNA Laboratory. Overall, we genotyped 36 individuals from the North Island and 13 individuals from the South Island at 22 potential microsatellite loci (S1 Table). Out of the 22 primers we optimized, four were unusable. For primers TAWH49 and TAWH33, electropherogram peaks could not be scored as microsatellites and were discarded. Primers TAWH7 and TAWH139 were monomorphic, only showing peaks at 174 bp and 210 bp respectively, and this was observed in every individual that had amplification at these loci (N = 8 and 30 respectively), so these primers were also discarded. Primer TAWH46 had consistent non-target amplification at 124 bp in almost all individuals so this peak was excluded, but we observed clear peaks at 150 or 154 bp that were retained and scored as microsatellites. Ultimately, we retained 18 microsatellite loci for analysis. Table 1 includes the primer sequences, fluorescent labels, annealing temperatures for each microsatellite locus included in this study, expected product sizes, number of alleles detected, and multiplex groupings.

**Table 1.**
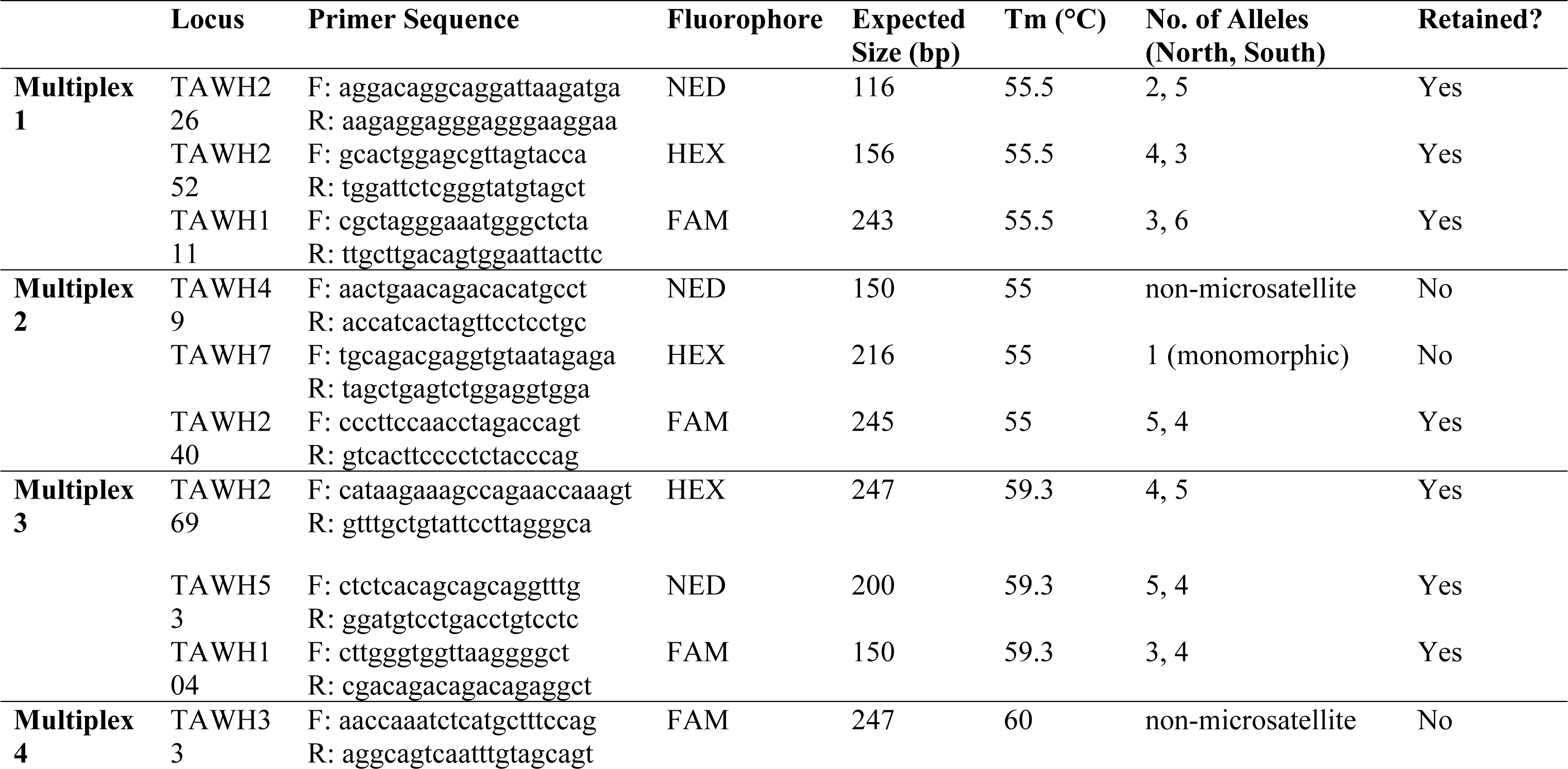
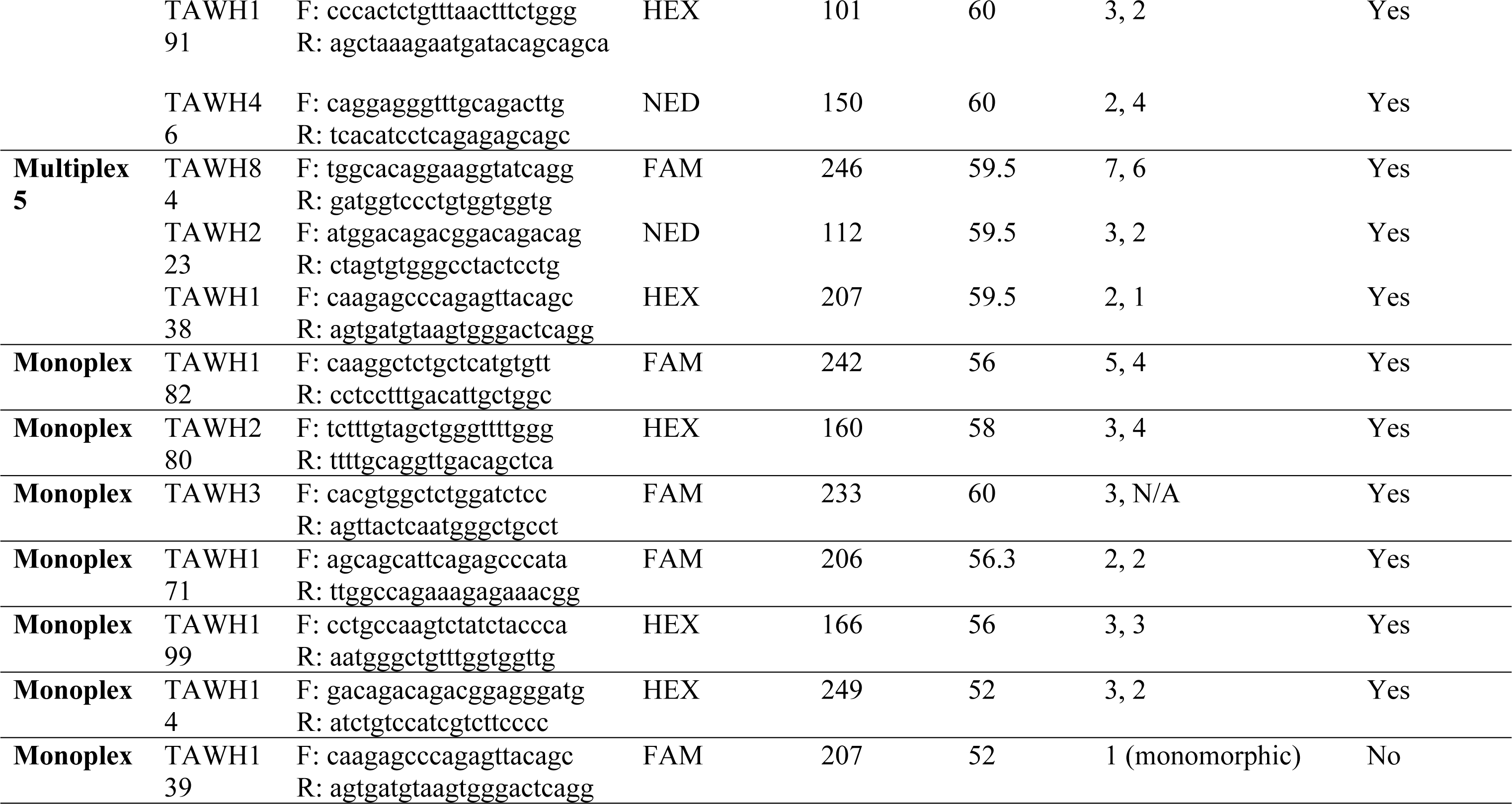
Details for each working multiplex, including primer name, sequence, expected product size, PCR recipe (with initial reagent concentrations), and thermocycling program. Thermocycling programs for multiplexes are as follows: Initial denaturation at 95°C for three min, followed by 30 cycles of denaturation at 95°C for 30 sec, annealing at X°C for 30 sec, and extension at 72°C for 45 sec, and a final extension at 72°C for 10 min.

### Statistical Analysis

To calculate heterozygosity, we scored electrograms using PeakScanner 2.0 using recommended methods in the Applied Biosystems DNA Fragment Analysis by Capillary Electrophoresis manual (2012). We used Arlequin v.3.2.2 (Excoffier and Lischer 2010) software for population genetic analyses, including calculations of Hardy-Weinberg Equilibrium (HWE), pairwise *F_ST_*, and Analyses of Molecular Variance (AMOVA). We used AMOVA to test for evidence of genetic structure within populations and to calculate conventional F-statistics; pairwise *F_ST_* was used to evaluate population structure between breeding groups within populations. A Mantel test was used to test for isolation-by-distance and was run with 999 permutations using the R package *vegan* (Oksanen et al. 2022). To correct for multiple tests, we applied a Bonferroni correction for AMOVA tests and pairwise *F_ST_* using the *stats* base package in R as described by Rice (1989). The significance of global *F_ST_* and F_IS_ between populations was tested by comparing bootstrap confidence interval values in a paired, two-tailed t-test. To visualize genetic differences between breeding groups and populations, we ran PCAs on raw allele counts using the R package *adegenet* (Jombart 2008), generating biplots of allelic similarity. Relatedness between individuals was investigated using COANCESTRY v.1.0.1.10 using a moment estimator (Wang 2002; Wang 2008). Heatmaps were generated using the *ggplot2* package in R (Wickham 2016). Finally, allelic diversity was determined using pairwise nucleotide diversity *pi* values calculated with Arlequin v.3.2.2 (Excoffier and Lischer 2010).

## Results

### Multiplexes

We successfully optimized five multiplexes, each with three primer sets, and six additional monoplexes (Table 1) for a total of 22 microsatellite loci. Each primer pair also amplified reliably in monoplex using the same PCR conditions as reported for multiplex reactions (S2 Table). Genotyping multiple individuals at each locus revealed allelic polymorphism in 18 microsatellites, 2 monomorphic microsatellites, and 2 unscorable microsatellites (Table 1). TAWH49 and TAWH33 may produce scorable results in monoplex, even though we did not obtain useable data in multiplex.

### Hardy-Weinberg Equilibrium (HWE) Tests

For all 18 polymorphic microsatellite loci and 6 breeding groups genotyped there were only 3 loci (TAWH53, TAWH269, TAWH223) that departed significantly from HWE expectations. Each departure was for only one breeding group (S3 Table) and each was due to a deficiency of observed heterozygotes. Within the North Island population, the HAY1N breeding group had four monomorphic loci and two loci that were out of HWE, with observed heterozygosity significantly lower than expected (S3 Table). The NPBS breeding group had six monomorphic loci and no loci deviating significantly from HWE. Lastly, RFSE and NPBE groups each had seven monomorphic loci and none departed significantly from HWE. The South Island population had two monomorphic loci and one locus that deviated from HWE due to lower-than-expected heterozygosity. Breeding groups W1 and W2 each had three monomorphic loci, two of which were the same loci. Group W1 had one locus that deviated significantly from HWE, while W2 had none (S3 Table).

### Population Structure and Relatedness

We found significant differentiation among breeding groups within the North Island population, with an *F_ST_* value of 0.24 (probability of non-departure from zero <0.01; Table 2A). Pairwise *F_ST_* scores among North Island breeding groups were all significant using Bonferroni corrected *p* values (Table 3). *F_ST_* values contrasting dyads of breeding groups from North Island were not significantly correlated with the distances between the nests of those groups in the dyads (Mantel test, p = 0.71).

**Table 2.** North Island (Tāwharanui) and South Island (Otokia) whole population AMOVA tests. Global AMOVA results are expressed as a weighted average over polymorphic loci. P-values adjusted using Bonferroni correction. Significance was tested using randomization tests with 15 000 permutations and 95% confidence intervals (CI) are indicated.

| A) North Island (N = 27 adults) |  |  |  |
| --- | --- | --- | --- |
| Source of variation | Sum of squares | Variance components | Percentage genetic variation |
| Among breeding groups (N = 4) | 79.8 | 1.44 | 23.8 |
| Among individuals within breeding groups | 161.3 | 2.11 | 34.9 |
| Within individuals | 70.0 | 2.50 | 41.3 |
| $F_{IS} = 0.458$ CI: 0.329 – 0.595 | | | |
| $F_{ST} = 0.238$ CI: 0.141 – 0.342 | | | |
| $F_{IT} = 0.587$ CI: 0.479 – 0.700 | | | |
| $F_{IS}$: $p = 0.000$ | | | |

| $F_{ST}$: $p = 0.000$ | | | |
| --- | --- | --- | --- |
| $F_{IT}$: $p = 0.000$ | | | |
| B) South Island (N = 8 adults) |  |  |  |
| Source of variation | Sum of squares | Variance components | Percentage genetic variation |
| Among breeding groups (N = 2) | 5.25 | -0.24 | -4.66 |
| Among individuals within breeding groups | 42.90 | 1.82 | 35.84 |
| Within individuals | 28.00 | 3.50 | 68.80 |
| $F_{IS} = 0.342$ CI: 0.148 – 0.538 | | | |
| $F_{ST} = -0.047$ CI: -0.107 – 0.014 | | | |
| $F_{IT} = 0.312$ CI: 0.131 – 0.502 | | | |
| $F_{IS}$: $p = 0.000$ | | | |
| $F_{ST}$: $p = 1.000$ | | | |
| $F_{IT}$: $p = 0.000$ | | | |

**Table 3.** Pairwise *F_ST_*for comparisons between and within North Island (Tāwharanui; HAY1N, NPBS, NPBE, RFSE) & South Island (Otokia; W1, W2) breeding groups. P-values adjusted using Bonferroni correction. Green: significant at alpha = 0.05; red: non-significant. Departure from the null hypothesis of F_ST_ being equal to zero was assessed using 15 000 permutations. Only comparisons between breeding groups in the same location are reported.

|  | HAY1N | NPBS | NPBE | RFSE | W1 | W2 |
| --- | --- | --- | --- | --- | --- | --- |
| HAY1N | 0.00 |  |  |  |  |  |
| NPBS | $F_{ST} = 0.191$<br>$p = 0.006$ | 0.00 | | | | |
| NPBE | $F_{ST} = 0.306$<br>$p = 0.002$ | $F_{ST} = 0.326$<br>$p = 0.006$ | 0.00 | | | |
| RFSE | $F_{ST} = 0.297$<br>$p = 0.018$ | $F_{ST} = 0.294$<br>$p = 0.006$ | $F_{ST} = 0.144$<br>$p = 0.024$ | 0.00 | | |
| W1 |  |  |  |  | 0.00 |  |
| W2 | | | | | $F_{ST} = 0.005$<br>$p = 0.915$ | 0.00 |

No population subdivision was detected between the two South Island breeding groups (*F_ST_* = 0; p = 1.00; Table 2B). Pairwise *F_ST_* scores between South Island breeding groups was 0.005, which was not significantly different from zero (p = 0.92; Table 3). To compare F-statistics for the two populations, we used 95% confidence intervals. The *F_ST_* and *F_IS_*values at North Island were both significantly higher than South Island (p <0.0001, Table 2A&B). Unexpectedly, F_is_ values for the South Island samples were statistically significant (Table 2B).

A PCA contrasting North and South Island populations indicated they cluster separately, with little to no overlap (Fig 2). Within North Island, HAY1N clusters the farthest from other breeding groups, with NPBE and NPBS clustering closer to RFSE. The two South Island breeding groups overlap with each other.

**Fig 2.**
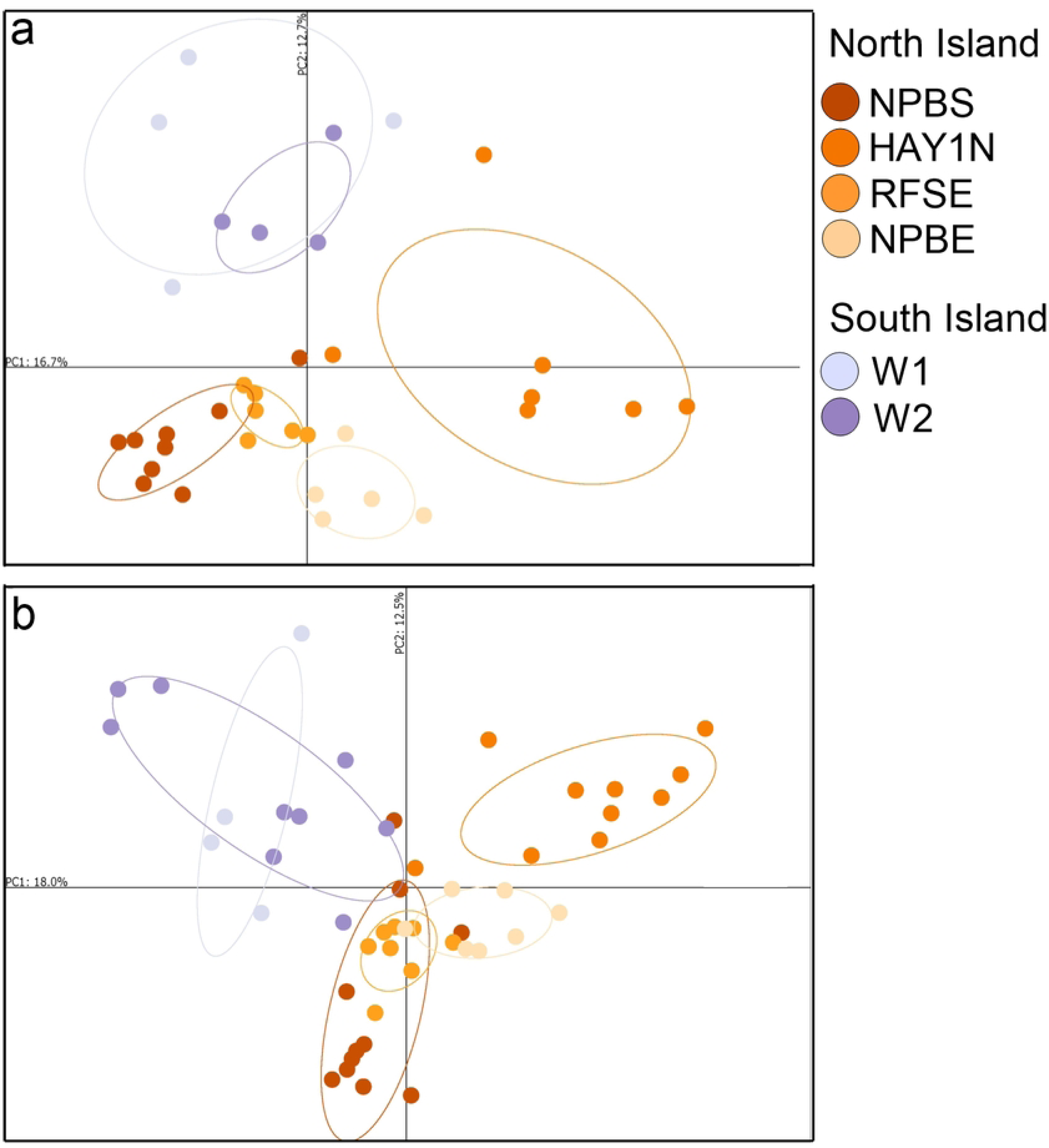
PCA of breeding groups from North Island (oranges) and South Island (purples) populations a) including offspring b) excluding offspring. Ellipses of inertia show where the majority of individuals cluster for each group. Figure generated using the *adegenet* package in R (Jombart 2008) and legends were generated in GraphPad Prism (version 10.0.0. for Windows, GraphPad Software, Boston, Massachusetts USA, www.graphpad.com).

As expected, relatedness within breeding groups was higher at the North Island population than the South Island population (Fig 3). Individuals in North Island breeding groups had higher coefficients of relatedness (*r* = 0.5 – 1.0), whereas between breeding groups, relatedness was comparatively lower (Fig 3). South Island breeding groups did not have a high coefficient of relatedness, apart from one dyad with *r* = 0.5 (see S4 Table for dyad relatedness values and 95% confidence intervals).

**Fig 3.**
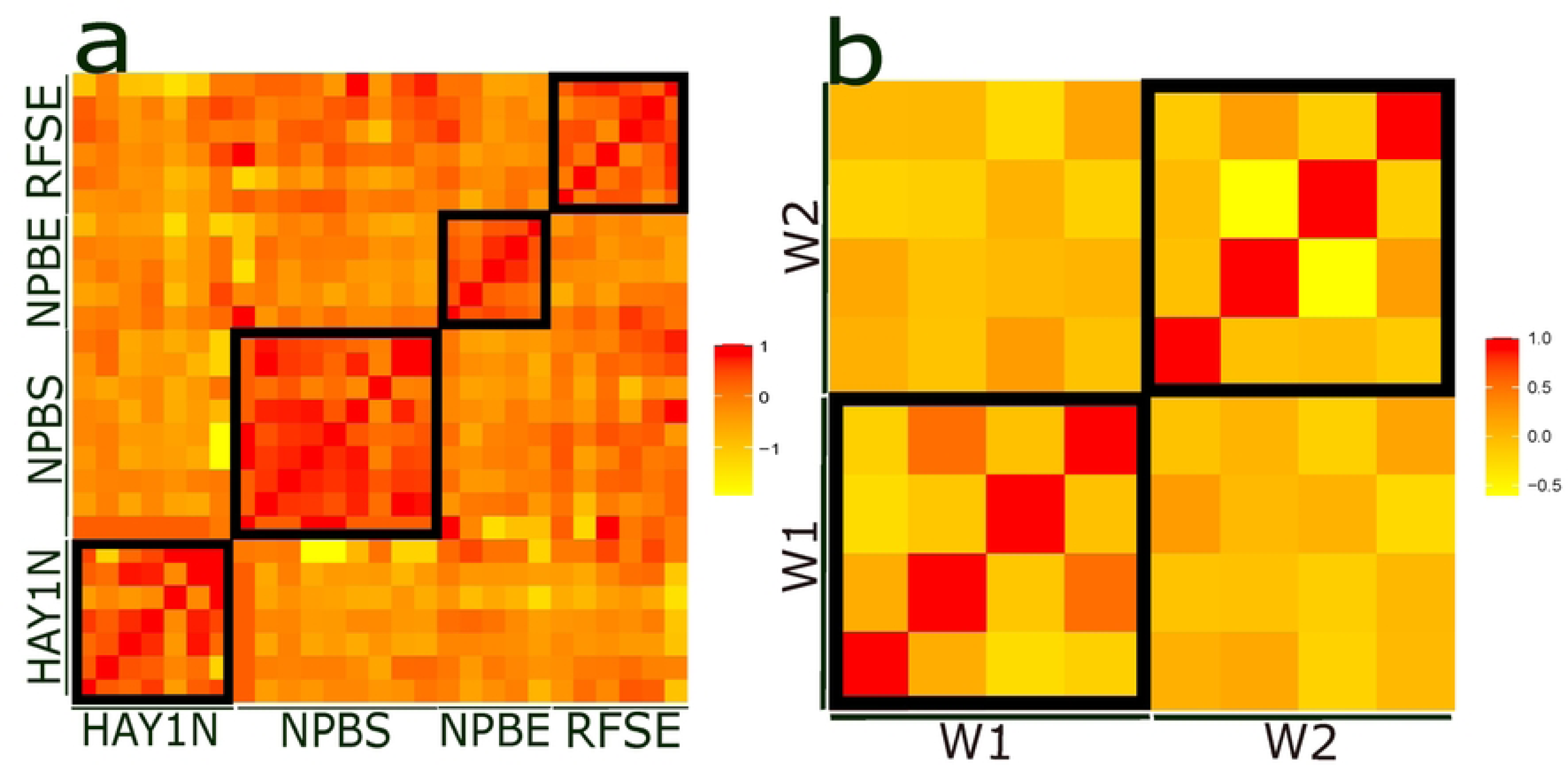
Heat maps depicting dyadic coefficients of relatedness. a) North Island (Tāwharanui) breeding groups b) South Island (Otokia) breeding groups. Bold outlines indicate within breeding group comparisons.

### Allelic Diversity

For North Island and South Island breeding groups, *pi* was similar and comparisons between breeding groups in these locations revealed no significant differences in allelic diversity (Table 4).

**Table 4.** Within breeding group *pi* values from each location.

| North Island |  |  |  | South Island |  |
| --- | --- | --- | --- | --- | --- |
| HAY1N | NPBS | NPBE | RFSE | W1 | W2 |
| 9.90 | 9.10 | 7.73 | 8.64 | 9.86 | 10.39 |

## Discussion

We successfully identified 18 polymorphic microsatellite markers, designed PCR primers that amplified their alleles, and used these data to test for population genetic verification of patterns that were discovered based on demographic and behavioural data from breeding groups that occupied year-round territories at Tāwharanui and Shakespear regional parks in the North Island. The genetic results reported here represent a first look that will be followed by a study of over 1000 adult blood samples collected since 2008 from > 60 North Island (Tāwharanui) groups. The main goal of this research was to establish a set of genetic tools for the study of parentage and kinship, as well as the genetic consequences of philopatry and year-round territoriality. These 18 newly developed polymorphic microsatellites, along with other markers to be developed, will allow us to study the consequences of inbreeding on individuals, breeding groups, and populations that are currently healthy and readily available for study. Three loci were significantly out of HWE in one of the 6 breeding groups (S3 Table). These three loci may have null alleles, which we will explore in the future.

We used these new microsatellite loci to test for genetic structure in four breeding groups from the Tāwharanui population on the North Island of New Zealand and two groups from the Otokia population on the South Island. While other studies have shown genetic structuring of cooperative breeding groups (see below), ours is the first study of a species with co-breeding males and females. This system, structured by breeding groups in the North Island population, promotes frequent inbreeding unless strong inbreeding avoidance mechanisms are in place.

### Genetic Structure, Group Kinship, and Inbreeding

Breeding groups in the North Island are genetically differentiated from each other based on AMOVA analysis (Table 2) and significant positive *F_ST_*values for all pairwise comparisons (Table 3). The mostly nonoverlapping grouping of individuals in each breeding group in the North Island, but not South Island, populations in a principal component analysis (Fig 2) is consistent with the *F_ST_* results. This was expected given field observations that neither offspring, nor adults, relocate between territories within Tāwharanui park in the North Island (*Pers. Obs.* J.S.Q.). Over the years of study since 2008, we found one banded individual, discovered by chance, that had migrated from our North Island study site. That bird dispersed over water (Kauwa Bay) approximately 10.5 km to Scandrett Regional Park. We have not yet systematically studied migration away from the North Island study site.

Inbreeding coefficients (*F_IS_* values) partially supported our prediction that birds from North Island are more inbred than those from South Island. The North Island population *F_IS_* values were significantly higher compared to those for the South Island population, supporting our prediction (Table 2). However, the *F_IS_*values in South Island were highly statistically significant, (Table 2B).

Microsatellite genotyping is prone to null alleles, which occur when an allele fails to amplify during PCR, sometimes due to mutations not allowing for primers to bind (Dakin and Avise 2004). This leads to an underestimation of heterozygosity and a consequent overestimation of homozygosity. It is possible that null alleles caused an overestimate of inbreeding (*F_IS_*) at both sites (S3 Table). For example, in the North Island, we observed 4 – 7 monomorphic loci in samples from each breeding group. In the South Island, we observed 3 monomorphic loci in each breeding group, but did not expect inbreeding. Null alleles may have caused an overestimate of homozygosity in our samples, leading to a less precise estimation of inbreeding (Chapuis and Estoup 2007; Waples 2018), however we do not expect that the significant difference between sites is in error. We will explore the possible null alleles using ML-NullFreq_frequency and other software to make sure whether they are real, and if so, how many actual null alleles there were (Dobrowski et al. 2015).

We predicted that pūkeko from the North Island would show signals of higher kinship resulting from strong philopatry and more frequent kin-matings within breeding groups. Indeed, we found that the North Island population was structured by breeding group differentiation, based on significant *F_ST_* values. Additionally, high coefficients of relatedness were common within, and much less so between, breeding groups in the North Island and not South Island (Fig 3). This pattern matches demographic and behavioural observations (Jamieson 1997; *Pers. Obs*. J.S.Q.). It appears that high levels of inbreeding in groups on year-round territories has led to a structured North Island population with frequent close kinships among group members (Wang and Shete 2011). We do not have evidence for genetic structuring in the South Island at Otokia, nor did we expect to find it there, but we acknowledge that our present analysis is based on only two breeding groups.

High relatedness coefficients within each of the four North Island breeding groups (Fig 3) suggest heightened potential for inbreeding. Having multiple male and female breeders in groups of polygyandrous pūkeko makes this example distinct from other cooperative breeding systems in terms of genetic structure. Several other cooperative breeding bird species show similar patterns of genetic structure based on social groups (e.g., apostle bird, *Struthidea cinerea*, Woxvold et al. 2006; white-browed sparrow, *Plocepasser mahali*, Harrison et al. 2014; white-winged chough, *Corcorax melanorhamphos,* Leon et al. 2021; bell miner, *Manorina melanophrys*, Painter et al. 2009). Another cooperative breeding species (superb fairy wren, *Malurus cyaneus*) does not show similar genetic structure between groups (Double et al. 2005), but this species has a very complex social system that includes common extrapair paternity outside the social group (Mulder et al. 1994).

We hypothesized that the South Island population may have arisen due to a range expansion from the North Island, so we predicted South Island birds would have lower allelic diversity than the North Island due to this founder effect. Pairwise comparisons of allelic diversity (*pi*) revealed no difference between North Island and South Island breeding groups. However, a limitation of our study that may at least partly explain this result is the small sample size we used for these pilot analyses. We only genotyped individuals from two South Island and four North Island breeding groups, which may not reflect the allelic diversity of the whole population. Follow-up studies will improve these tests by expanding sample sizes for both populations.

## Conclusions

We conducted this pilot study to identify microsatellites, design new primers to amplify them, and test them on samples of wild caught pūkeko. We successfully designed five multiplexes, each containing three primer pairs, and 18 microsatellites were individually assayed. In total, we had 18 polymorphic loci, two monomorphic, and two unscorable microsatellite loci for pūkeko. One multiplex contained two loci that were unscorable in multiplex, but that may be scorable in monoplex, so we will explore the utility of these loci further in future studies. Using the 18 polymorphic loci, we estimated heterozygosity and inbreeding coefficients within and between breeding groups and between the North and South Island study populations. These robust protocols will be used by future researchers engaged in pūkeko population genetics studies and will also facilitate future investigations of kinship and parentage analyses in pūkeko.

Results from this study begin to explore differences between North and South Island New Zealand pūkeko populations using microsatellites by determining patterns of heterozygosity and inbreeding coefficients in a North Island pūkeko population with high levels of kin-mating compared with an outbred South Island population. Our analyses suggest that the North Island population lives in a structured population of inbred kin groups living and breeding on year-round territories (Craig and Jameison 1990), while the South Island population lives in a non-structured outbred population living on seasonal territories and mixing in the non-breeding season (Jamieson 1997). Our pilot study presents preliminary genetic analyses that may be useful to study pūkeko as a model for wild inbreeding populations that will help us understand the fitness impacts of inbreeding in declining endangered populations and how inbreeding depression and loss of heterozygosity contributes to fitness decline.

## Acknowledgments

We thank Matt Maitland, Staff, Rangers, and TOSSI (Tāwharanui Open Sanctuary Society Inc.) members for their kindness and support during our field work at Tāwharanui Regional Park. Dr. Cody Dey, assisted J.S.Q. in collecting demographic and behavioural data in 2010 for the general pukeko project.

## Supporting Information

**S1 Table. Genotyped individuals.** Samples included in analyses.

**S2 Table. PCR details**. Primer sequences, fluorophores, expected size and melting temperature and conditions for PCR of multiplexes and singleplexes.

**S3 Table. Hardy-Weinberg equilibrium tests for North Island (Tāwharanui) and South Island (Otokia) breeding groups.** Tests have 1 000 000 steps in Markov chain and 100 000 dememorization steps. Rows with red text correspond to p <0.05. N = number of individuals, #ind = number of adults genotyped at a given microsatellite locus.

**S4 Table. Dyadic relatedness**. Wang relatedness values and 95% confidence intervals.

**S1 File. Ethanol Precipitation.** Ethanol precipitation (‘cleanup’ or ‘desalting’) of PCR products to be sent to Trent for genotyping on the ABI 3730

## Notes

### Competing Interest Statement

The authors have declared no competing interest.

